# scSLAM-seq reveals core features of transcription dynamics in single cells

**DOI:** 10.1101/486852

**Authors:** Florian Erhard, Marisa A.P. Baptista, Tobias Krammer, Thomas Hennig, Marius Lange, Panagiota Arampatzi, Christopher Jürges, Fabian J. Theis, Antoine-Emmanuel Saliba, Lars Dölken

## Abstract

Current single-cell RNA sequencing approaches gives a snapshot of a cellular phenotype but convey no information on the temporal dynamics of transcription. Moreover, the stochastic nature of transcription at molecular level is not recovered. Here, we present single-cell SLAM-seq (scSLAM-seq), which integrates metabolic RNA labeling, biochemical nucleoside conversion and single-cell RNA-seq to directly measure total transcript levels and transcriptional activity by differentiating newly synthesized from pre-existing RNA for thousands of genes per single cell. scSLAM-seq recovers the earliest virus-induced changes in cytomegalovirus infection and reveals a so far hidden phase of viral gene expression comprising promiscuous transcription of all kinetic classes. It depicts the stochastic nature of transcription and demonstrates extensive gene-specific differences. These range from stable transcription rates to on-off dynamics which coincide with gene-/promoter-intrinsic features (Tbp-TATA-box interactions and DNA methylation). Gene but not cell-specific features thus explain the heterogeneity in transcriptomes between individual cells and the transcriptional response to perturbations.

## Main

Regulation of gene expression is a fine-tuned process, which allows cells to maintain homeostasis and respond to changing environmental conditions. Single-cell RNA sequencing (scRNA-seq) allows to quantify transcript levels for thousands of genes in individual cells. This revealed that intercellular heterogeneity plays an important role in phenotype variability in health and disease ^1–4^. However, current scRNA-seq approaches only quantify total cellular RNA profiles rather than transcriptional activities. More importantly, the RNA profile of each individual cell can only be analyzed once at a given time point. This precludes direct monitoring of transcriptional changes in individual cells due to perturbations. Accordingly, changes can only be inferred from differences in transcript levels between distinct cell populations or conditions ^5^. Furthermore, gene expression is a stochastic process, with intrinsic and extrinsic noise in transcription and translation leading to intercellular heterogeneity in mRNA and protein levels ^6^. These inherent characteristics of transcription are not resolved by current scRNA-seq approaches.

### Single-cell SLAM-seq

Here, we introduce single-cell SLAM-seq (scSLAM-seq) to directly quantify transcriptional activity in single cells and resolve the involved regulatory elements. We validate scSLAM-seq by profiling changes occurring after challenge with a virus. scSLAM-seq is based on thiol(SH)-linked alkylation for the metabolic sequencing of RNA (SLAM-seq) ^7,8^, which involves a brief exposure of cells to the nucleoside analog 4-thiouridine (4sU). 4sU is incorporated into RNA during transcription and converted to a cytosine analog using iodoacetamide (IAA) prior to RNA-seq. Sequencing reads originating from “new” RNA, which has been transcribed during the time of 4sU labeling, can be identified within the pool of total RNA reads based on characteristic U- to C-conversions. We applied scSLAM-seq to decipher intercellular heterogeneity in the initial cellular response to lytic murine cytomegalovirus (CMV) infection at a time of infection (2hrs) when the bulk of transcriptional regulation still remains undetectable in total cellular RNA^9^. After improving and optimizing the SLAM-seq protocol for single-cells sequencing (Fig. 1a), we performed scSLAM-seq on 107 single murine fibroblast cells and in parallel analyzed global transcriptional changes of a matched larger (1×10^5^) population of cells using SLAM-seq. After filtering out ‘low quality’ cells (Extended Data Fig. 1a), the remaining (49 CMV-infected, 45 mock-infected) showed all characteristics of high-quality scRNA-seq and SLAM-seq libraries (>2,500 quantified genes, characteristic distributions of viral/cellular/spike-in/mitochondrial RNA (Extended Data Fig. 1a,b); U- to C-conversion rates between 4 and 6% (Fig. 1b,c and Extended Data Fig. 1c)). Overall, we found 4sU incorporation to be both efficient and uniform at single cell level and comparable to SLAM-seq performed on large populations of cells.

**Figure 1.**
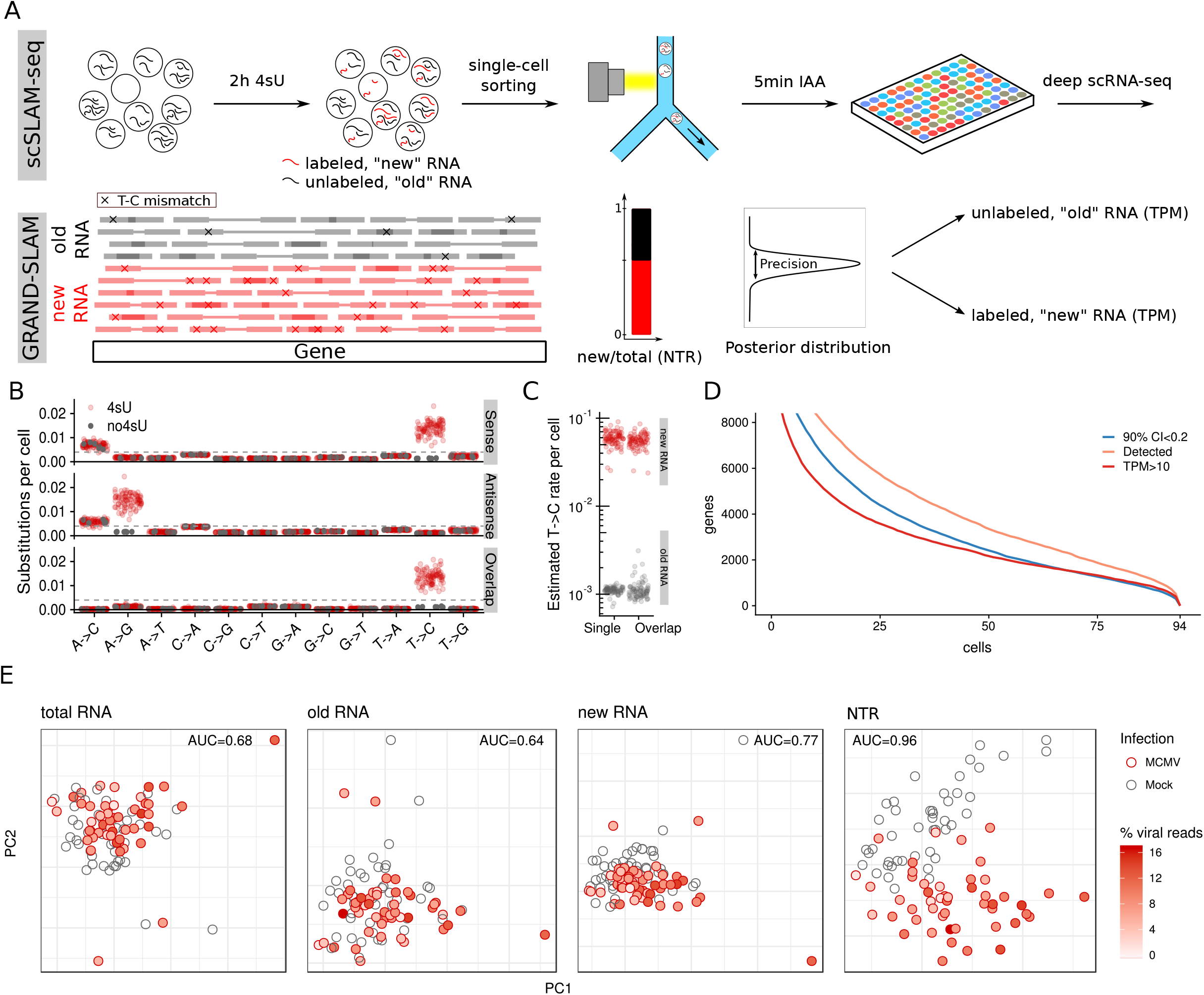
scSLAM-seq resolves transcriptional activity at single-cell level. **a**, Overview of scSLAM-seq and GRAND-SLAM. Nascent transcripts are labeled by 4sU for 2hrs. Following single-cell sorting and RNA isolation, 4sU is converted into a cytosine analogue by iodoacetamide (IAA) and SMART-seq libraries are prepared and sequenced. GRAND-SLAM identifies thymine to cytosine mismatches and estimates both the ratio of newly transcribed to total RNA (NTR) and expression estimates of “old” and “new” RNA (TPM: transcript per millions). **b**, Rates of nucleotide substitutions demonstrate efficient conversion rates in 4sU-treated single cells (4sU) compared to 4sU-naïve cells (no4sU). This was true for reads originating from both cDNA strands (sense and antisense) as well as overlapping parts of the paired-end sequencing (overlap). **c**, Estimated T- to C-substitution rates in new and old RNA determined by GRAND-SLAM in non-overlapping (single) and overlapping parts of the paired-end sequencing. **d**, Number of genes for which the NTR could be quantified with high precision (90% credible interval (CI) <0.2) compared to the detected genes and reliably detected genes (TPM>10). **e**, Principal component analysis (PCA) on highly variable cellular genes separates infected and mock cells based on the NTR only (AUC: Area under the receiver operating characteristics curve; PC: principal component).

Previous nucleoside conversion approaches utilized the number of sequencing reads with U- to C-conversions as a semi-quantitative measure for new RNA ^7,8,10^. However, due to the low 4sU incorporation rates of about 1 in 50-200 nucleotides, a substantial fraction (up to >50%) of all SLAM-seq reads that originate from new RNA do not contain U- to C-conversions. Moreover, reads from unlabeled (“old”) RNA also contain U- to C-substitutions due to sequencing errors, PCR errors, SNPs, RNA editing or transcription errors. To overcome these confounding factors, we previously developed *Globally refined analysis of newly transcribed RNA and decay rates using SLAM-seq* (GRAND-SLAM), which employs statistical methods to compute the ratio of newly transcribed to total RNA (NTR) in a fully quantitative manner including credible intervals ^11^ (Fig. 1a-c). Here, we report on GRAND-SLAM 2.0 for parallel analysis of hundreds of SLAM-seq libraries derived from single cells. Quantification accuracy is further improved by analyzing long reads in paired-end mode (See Methods), where partially overlapping sequencing reads allow to reliably distinguish 4sU conversions from sequencing errors (Fig. 1b, Extended Data Fig. 1c). Thereby, NTRs including confidence scores thereof are determined for thousands of genes per single cell. The number of genes for which the NTR could be estimated with high precision approached the overall sensitivity of single-cell sequencing (Fig. 1d). The expression estimates for total, new and old RNA after pooling the single-cell data showed a strong correlation (R between 0.73 and 0.77) with the bulk expression data (Extended Data Fig. 1d). This demonstrates that scSLAM-seq combined with GRAND-SLAM 2.0 accurately resolves transcriptional activity for thousands of genes in single cells. It allows us to analyze the transcriptional profiles prior to, during and at the end of the window of 4sU-labeling thereby introducing a temporal scale to scRNAseq.

### CMV transcriptional regulation

Short-term changes in transcriptional activity that occur within a two-hour window largely remain undetectable in total RNA levels due to RNA half-lives in the order of hours for most mammalian transcripts ^12^. At single cell level, they are further obscured by intercellular differences in transcript levels. Accordingly, unbiased principal component analysis (PCA) on highly variable genes (Methods, Extended Data Fig. 1e) could not separate CMV-infected cells from mock-infected cells for either total RNA or old cellular RNA, and only barely for new RNA (Fig. 1f) when viral reads were excluded (Extended Data Fig. 1f-g). The heterogeneity in gene expression between individual cells thus exceeds the virus-induced alterations. The NTR places transcriptional activity in relation to the RNA levels that were present prior to infection. PCA on the NTR separated uninfected from infected cells with high precision. Moreover, there was a clear positive correlation for the PCA distance of an infected cell from the uninfected cells with the extent of viral gene expression (R=0.46, p=0.00095; Extended Data Fig. 1h). Based on RNA half-lives of >13,000 genes from our cells (Supplementary Table 1, Extended Data Fig. 2a), we found that the poor discriminative power of total RNA reflects its poor temporal resolution for picking up transcriptional changes of genes with medium- to long-lived transcripts (t_½_ > 4hrs; see Extended Data Fig. 2b-d).

Cytomegalovirus gene expression occurs in a regulated cascade of immediate early, early and late gene expression. Late gene expression is dependent on viral DNA replication and in fibroblasts initiates around 12h of infection ^9^. In addition, herpesvirus virions deliver substantial amounts of RNA from all three kinetic classes into the infected cells, which masks the onset of viral gene expression when analyzing total cellular RNA ^13,14^. We previously reported on a peak of viral gene expression of all kinetic classes during the first 2hrs of CMV infection using 4sU-seq on bulk cells ^9^. The origin and nature of the respective transcription, however, remained elusive. scSLAM-seq now allowed us to assess this issue in individual CMV-infected cells. Consistent with the efficient initiation of productive infection, the majority of viral immediate early and early transcripts in all cells were new (Fig. 2). However, the same picture was also observed for the vast majority of late genes, which should not have been transcribed within the first two hours of infection. Interestingly, transcriptional activity of viral genes from all kinetic classes also represents one of the first events in herpes simplex virus 1 reactivation, termed “Phase I” ^15,16^. It is thought to reflect a state in which the previously repressed viral genomes become totally de-repressed. Our data indicate that the naked incoming CMV DNA in permissive cells similarly facilitates the expression of the majority of viral genes independent of their kinetic class. The breath of viral gene expression and our poly(dT) primer-based scRNA-seq approach strongly argue that the respective transcripts are properly processed and presumably translated. Only for a small set of viral genes, transcripts were devoid of U- to C-conversions. Interestingly, some of these were amongst the most abundant viral transcripts in the infected cells at 2hrs post infection. The respective transcripts thus represent virion-associated RNAs thereby providing a quantitative measure of the number of virus particles the respective cell has been infected with two hours ago. The dose of infection exhibited a strong positive correlation with the extent of *de novo* viral gene expression at single cell level (R^2^=0.66, Extended Data Fig. 3). This is consistent with the well-described role of many viral tegument proteins in overcoming intrinsic cellular antiviral resistance thereby augmenting the onset of productive infection ^17^.

**Figure 2.**
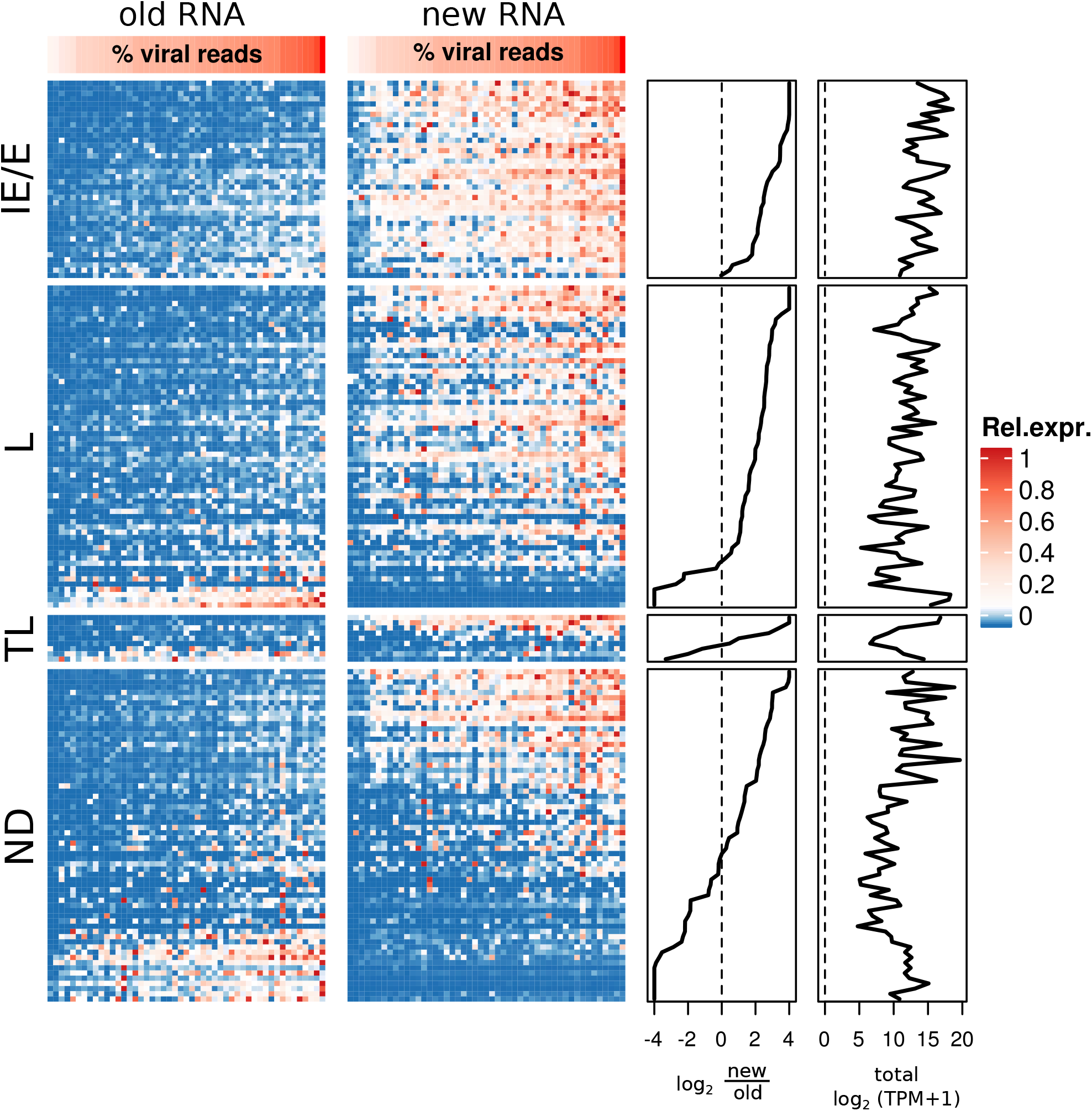
Pervasive viral gene expression of all kinetic classes and incoming RNA revealed by scSLAM-seq. Heatmaps showing the levels of old and new RNA relative to the maximal total level for each viral gene (rows) per CMV infected cell (columns). Cells are sorted according to the percentage of viral reads among all reads from the cell. Kinetic classes are indicated (I/E: immediate early and early, L: late, TL: true late, ND: not defined). The ratio of new to old RNA (log_2_ new/old) and total expression from pooled cells for each viral gene is depicted.

To assess the effects of CMV infection on cellular gene expression, we identified differentially expressed genes ^18,19^ between CMV-infected and mock-infected single cells in total, new and old RNA (Supplementary Table 2). The majority (>60%) of both downregulated (61/87) and upregulated genes (188/309) could only be uncovered by new RNA (Fig. 3a and Extended Data Fig. 4a-b). Consistent with results from population based experiments ^9^, this implies that the majority of differentially expressed genes are regulated at transcriptional level and not due to changes in RNA degradation. Changes in gene expression detected in large cell populations can either result from a global, and relatively mild change in transcription in all cells in the population (“Up” or “Down”) or in a selective change in the number of cells transcribing the respective gene (“Off” vs “On”). ScSLAM-seq now allows to determine which of these fundamentally different modes of regulation applies to each individual gene. Considering the strong intercellular differences in transcriptional activity (new RNA) even for highly expressed genes, our data demonstrate that CMV-induced transcriptional alterations within the first two hours of infection predominantly result from “Off - On” transcriptional dynamics (Fig. 3b and Fig. 3c-f for examples). This was particularly prominent for genes encoding for short-lived transcripts independent of their expression levels.

**Figure 3.**
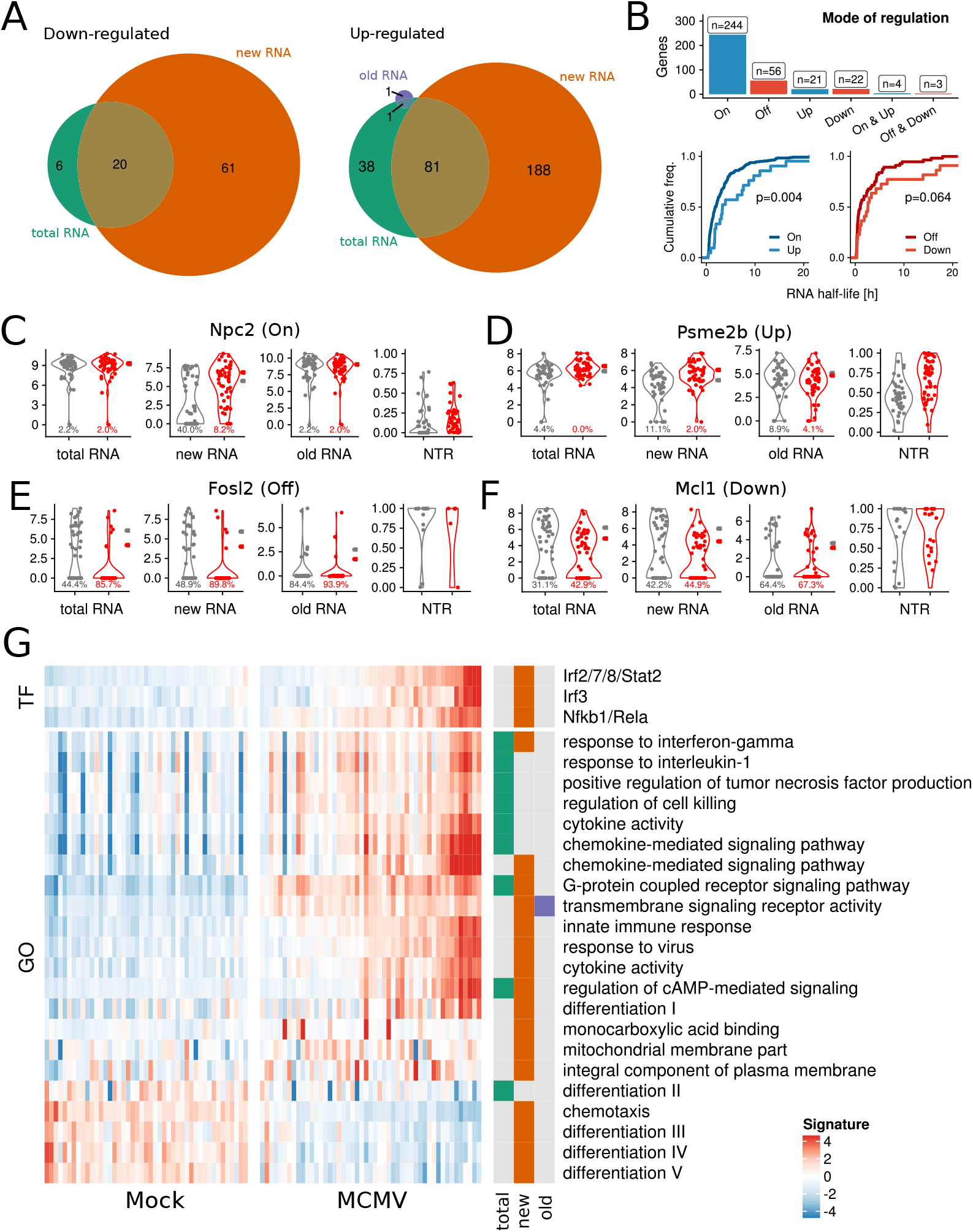
scSLAM-seq reveals the mode of gene regulation and differentially activated pathways in single cells. **a**, Pie charts representing up- and down-regulated genes upon CMV infection called using single-cell two-phase testing procedure (SC2P) (10% false discovery rate) and based on either total, new or old RNA. **b**, Top: Regulated genes in new RNA are classified by SC2P according to the mode of regulation (On/Up, Off/Down). Bottom: RNA half-life distributions of Up/Down genes are compared to On/Off genes, respectively (two-sided Wilcoxon rank-sum test). **c-f**, *Npc2, Psme2b, Fosl2, Mcl1* illustrate the different modes of regulation. Grey: mock, Red: CMV-infected cells. The percentage of cells not expressing a transcript is indicated below every violin plot. For total, new and old RNA, average expression levels are indicated to the right of every violin plot. The y axis indicates normalized read counts (log_2_). For new/total, the y axis indicates the NTR. **g**, Unbiased pathway and gene set overdispersion analysis (‘PAGODA’) revealed transcriptional signatures associated with mock- and CMV-infected cells. Upper and lower panels depict the identified transcription factors targets (TF) and gene ontology (GO) signatures, respectively. The fraction (total, new, old) in which each signature was found is indicated.

CMV infection is well described to induce a strong NF-κB and a type I interferon (IFN) response during the first 2hrs of infection ^9,20^. In which cells these responses are activated and how this relates to the progression of infection remained unclear. Gene set analysis ^21^ from predicted transcription factor targets ^22^ and gene ontology terms demonstrated pervasive NF-κB activation in the majority of the infected cells, whereas IFN activation was limited to about half of the infected cells (Fig. 3g, Supplementary Table 3 & 4). This is consistent with virion-mediated delivery of the viral tegument protein M45, which in turn directly activates NF-κB by degrading the NF-κB essential modulator (*Nemo*) ^20^. In contrast, IFN responses require the detection of pathogen-associated molecular patterns (PAMPS) ^23^ and are thus to a much greater extent influenced by intercellular heterogeneity ^24^. To perform a more focused analysis, we defined both NF-κB- and IFN-responsive gene sets specific to CMV infection based on the intersection of published data on NF-κB induction ^25^ and IFN treatment ^12^ with our bulk SLAM-seq data (Supplementary Table 5). The magnitudes of both the NF-κB (Fig. 4a,b) and IFN (Fig. 4c,d) responses varied markedly between individual cells but were highly correlated (Spearman’s ρ>0.52, p<3×10^−4^) with viral gene expression. The bulk of both the NF-κB- and IFN-inducible gene expression thus arises in the most strongly infected cells. In old RNA, expression levels of IFN-inducible genes were indistinguishable from those observed in uninfected cells (Fig. 4a). This is consistent with efficient differentiation of new from old RNA. However, expression levels of the NF-κB-responsive genes in old RNA of infected cells were significantly higher than in the uninfected cells (Fig. 4c). Following the exposure of cells to 4sU, metabolic labeling takes a few minutes to initiate. It subsequently increases in efficiency over time ^26^. We conclude that the induction of NF-κB signaling by the viral tegument M45 occurs more rapidly than the virus-induced activation of the IFN response.

**Figure 4.**
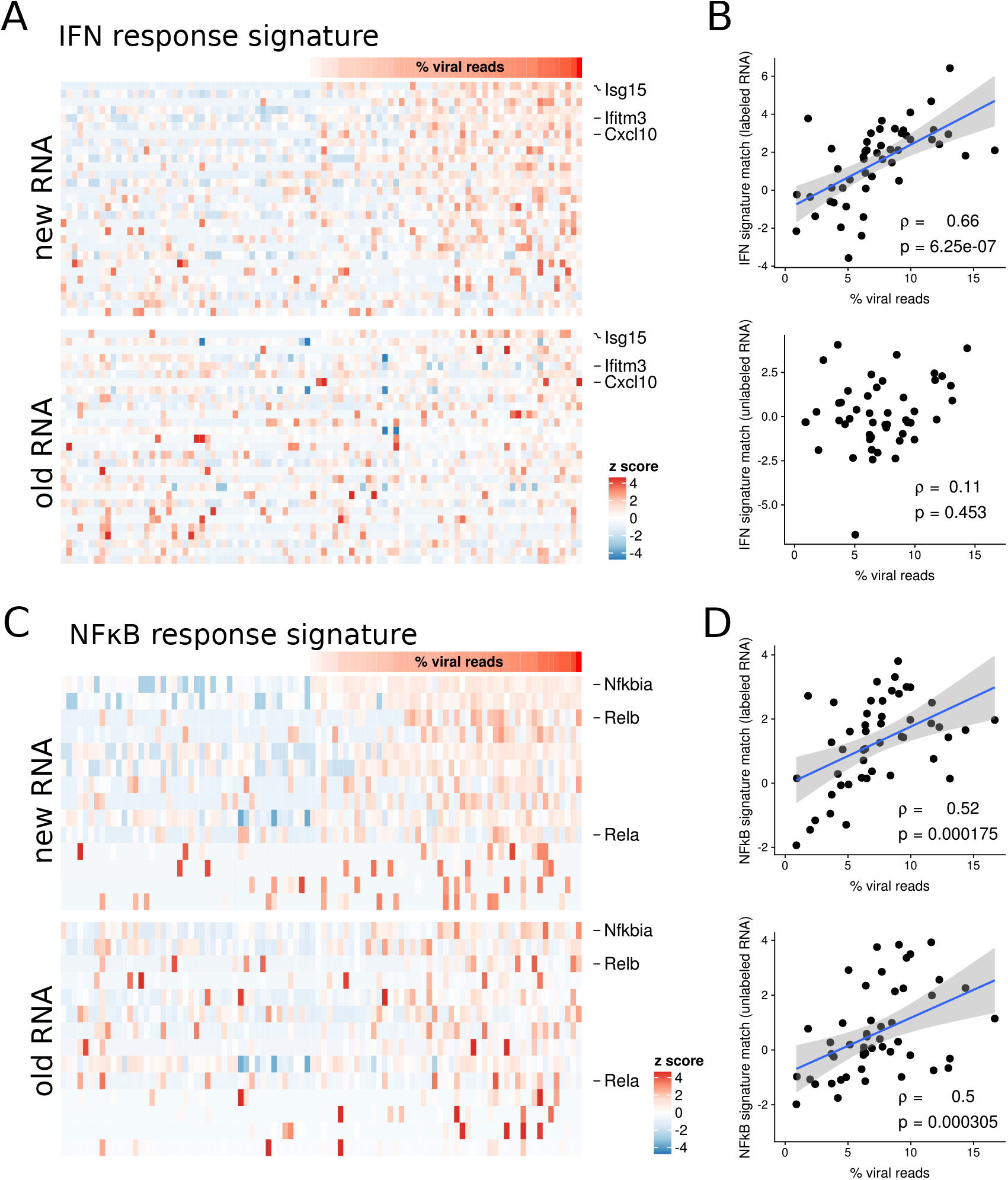
scSLAM-seq relates IFN and NF-κB responses to productive infection and outcome. **a**, Standardized expression levels of interferon (IFN)-responsive genes are depicted for new and old RNA. Cells are ordered according to the total percentage of viral reads. **b**, The IFN response signature score for each cell is plotted against its viral RNA content. The linear regression fit and its 95% confidence interval are indicated. **c-d**, Standardized NF-κB-responsive gene expression levels and signature scores are depicted as for IFN.

### Dichotomous transcription

For many cellular genes in the uninfected cells, we observed striking “On”-“Off” patterns in the RNA transcribed within the 2hrs of labeling. For instance, for both the master regulator Hypoxia Inducible Factor 1 Alpha Subunit (*Hif1a*) and Autophagy Related 12 (*Atg12*) all transcripts in individual cells were either new or old (Fig. 5a). In contrast, other genes such as Ferredoxin 2 (*Fdx1l*) and WD Repeat Domain (*Wdr89*) exhibited much less or no variability, respectively (Fig. 5a). Extreme cases of dichotomy such as Hif1a or Atg12 might be due to singular mRNAs captured from single cells, however in general the observed dichotomy was not a consequence of artifacts due to the minute amount of RNA in single-cell sequencing (Extended Data Fig. 5a). Moreover, it did not result from cell cycle-dependent differences (Extended Data Fig. 5b,c), but was correlated with RNA half-life (Extended Data Fig. 5d).

**Figure 5.**
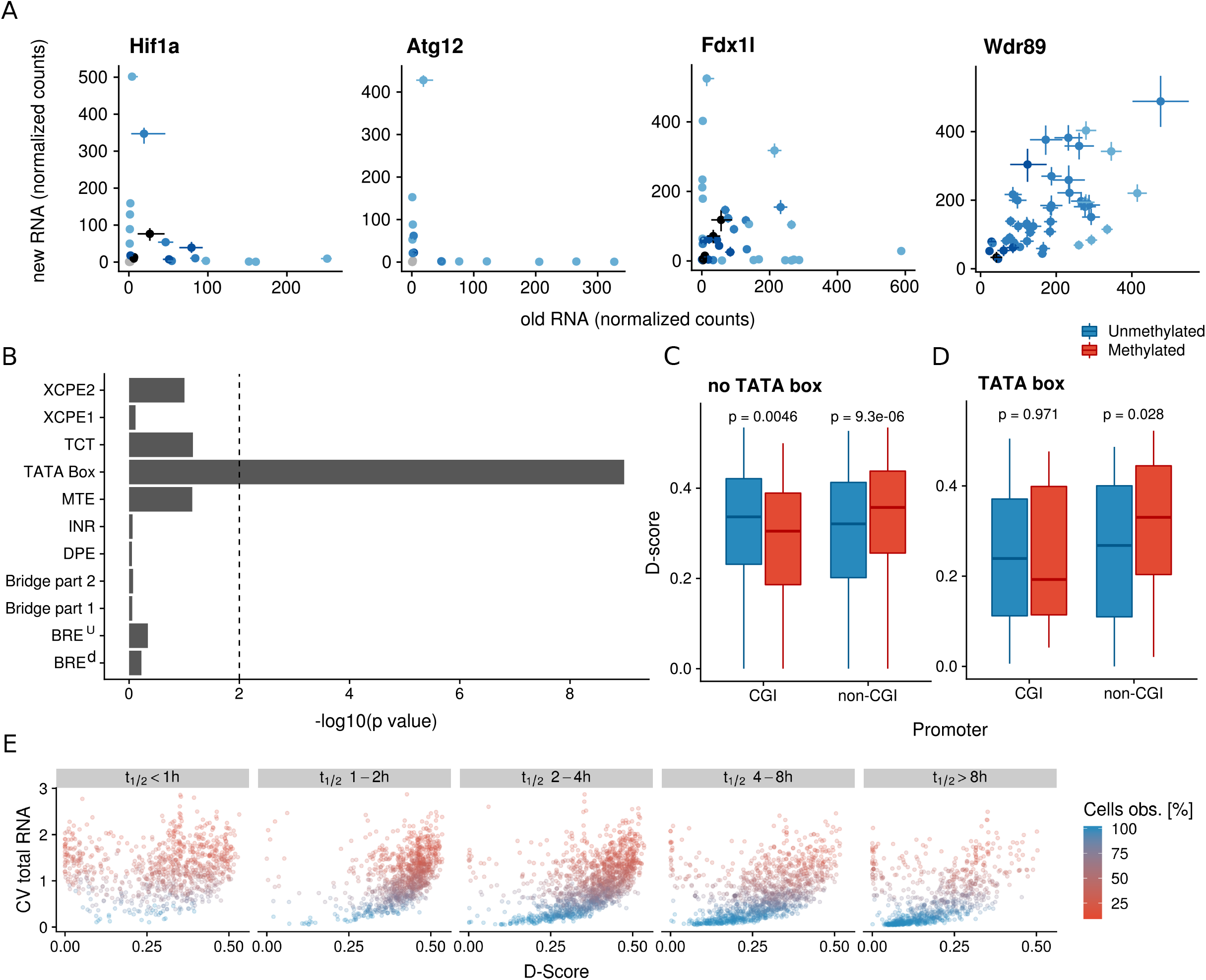
scSLAM-seq reveals dichotomous transcription and core features of heterogeneity in transcription. **a**, Examples of dichotomous (*Hif1a, Atg12*), intermediate (*Fdx1l*) and stable (*Wdr89*) transcriptional activity. Shown are the normalized read counts of old and new RNA for each cell with associated 90% credible intervals. **b**, Promoter structure analysis reveal TATA boxes to be highly enriched in promoters of non-dichotomous genes. Log-transformed p values are indicated (two-sided Wilcoxon rank-sum test). **c**, Dichotomy score (D-score) distributions for CpG island (CGI) promoters and non-CpG island (non-CGI) promoters stratified by DNA methylation status in bisulfite sequencing experiments are shown. Only promoters without TATA boxes were considered. P values for differences of scores in strata are indicated (two-sided Wilcoxon rank-sum test). **d**, The same as in **c** but considering TATA box promoters only. **e**, D-scores for each gene are scattered against the coefficient of variation (CV) of total RNA across cells. Genes were stratified according to their RNA half-life (t_½_). The percentage of non-drop-out cells is indicated. (XCPE1 & 2 = X core promoter element 1 and 2, TCT = polypyrimidine initiator element, MTE = Motif 10 element, INR = Initiator element, DPE = Downstream promoter element, BRE^u^ = TFIIB recognition element upstream of TATA box, BRE^d^ = TFIIB recognition element downstream of TATA box).

Transcriptional activity at single cell level is not a continuous process. Single-molecule imaging rather demonstrated intermittent burst of transcription separated by minutes to hours of transcriptional inactivity ^6,27–30^. To globally assess these effects within our scSLAM-seq data, we defined a gene-wise dichotomy score (D-score) as the standard-deviation of the NTR from all uninfected cells in which RNA of the respective gene was detected (Supplementary Table 6). Unbiased gene ontology overrepresentation analysis ^31^ revealed high D-scores to be associated with functional categories such as protein phosphorylation and ubiquitination (Supplementary Table 7). To assess the molecular principles of dichotomous transcription, we performed gene promoter analyses. This revealed six motifs to be significantly enriched for either low (TATA box motif) or high D-scores (CG-rich and purine-rich motifs; Extended Data Fig. 6; Supplementary Table 8). The former indicates that core promoter structure influences the frequency and duration of transcriptional inactivity. Indeed, promoter structure analysis revealed a highly significant (p<10^−8^) enrichment of correctly placed TATA boxes in promoters of genes with low D-score, but no association with any other core promoter motifs (Fig. 5b). This is consistent with TATA box promoters driving frequent and regularly spaced transcriptional bursts ^32^. In contrast, the observed CG-rich motifs could either correspond to binding sites of specific transcription factors that exhibit an oscillatory activation pattern, or reflect short stretches of CpG dinucleotides in the respective promoters. More than 50% of mammalian transcription initiates from promoters close to CpG islands (CGI promoters), which represent CpG-rich regions of dozens to hundreds of nucleotides ^33,34^. CpG island hypermethylation is an epigenetic control mechanism that is important for gene inactivation ^35^. Hypermethylation of CpG islands has been described in almost any kind of tumor and has been shown to be involved in silencing of many key pathways such as DNA repair, cell cycle and apoptosis ^36,37^. However, CpG methylation at non-CGI promoters may also control gene expression. We assessed the association of CpG islands and their methylation pattern with dichotomous transcription utilizing CGI promoter annotations downloaded from the UCSC genome browser and bisulfite sequencing data from mouse fibroblasts ^38^. DNA methylation of CGI promoters correlated with low D-scores whereas methylated non-CGI promoters exhibited high D-scores (Fig. 5c). Thus, DNA methylation rather than specific transcription factors coincides with dichotomous transcription and differentially affects CGI and non-CGI promoters. The same was also observed for TATA box-containing promoters (Fig. 5d). These findings are consistent with a model in which DNA methylation outside of CpG islands within gene promoters is involved in the transient regulation of promoter activity in the time-scale of hours by temporarily rendering gene promoters non-permissive. Interestingly, the resulting dichotomous transcription explains the heterogeneity in the transcriptomes of individual cells (Fig. 5e). The observed heterogeneity thus is not a cell-but a gene-specific phenomenon and highlights core promoter elements including TATA boxes and CpG composition as major factors defining intercellular heterogeneity in transcript levels.

## Discussion

Deciphering the real-time transcriptional dynamics with single-cell resolution is fundamental to understand how gene expression is orchestrated at molecular level. Here, we integrate 4sU metabolic labeling, biochemical nucleotide conversions and a new tailored bioinformatics approach to simultaneously visualize total transcript levels, transcriptional activity and to differentiate newly synthesized from pre-existing RNA. Thereby, the impact of intercellular heterogeneity on the response to a perturbation can be analyzed. This adds a temporal dimension to scRNA-seq, substantially increases the power of scRNA-seq to identify regulatory changes in gene expression and differentiates changes in RNA synthesis from alterations in RNA decay. Most recently, a multiplexed single-molecule in situ method, intron seqFISH was developed that allows imaging of >10,000 genes at their nascent transcription sites in single cells by mRNA and lncRNA seqFISH and immunofluorescence, thereby revealing fast dynamics and asynchronous oscillations ^39^. However, the approach is based on the assumption of rapid and stable splicing kinetics of all introns and is technically highly demanding. In contrast, scSLAM-seq directly measures newly synthesized RNA and is readily established by any single-cell sequencing facility only prolonging hands-on time during library preparation by ~30min.

scSLAM-seq depicts the stochastic nature of transcription genome-wide. Single molecule imaging studies on individual reporter genes visualized transcriptional bursts by convoys of polymerases followed by hours of transcriptional inactivity indicative of temporarily non-permissive promoters ^29,40^. Our approach reveals the concerted effects of transcriptional bursts and ongoing RNA decay on cellular transcript levels for thousands of cellular genes. In the most striking cases of dichotomous transcription, exemplified by *Hif1a* and *Atg12*, all mRNA molecules of the respective genes are degraded before the next burst occurs. Even for genes with a half-life of 1h or less, the respective promoters apparently do not initiate transcription for multiple hours. This provides strong evidence that the drop-outs frequently observed in single-cell sequencing even for genes highly expressed on the population average are not artifactual, but indeed correspond to prolonged periods of inactivity (off-state) of the respective gene promoters resulting in a near-complete loss of these mRNAs in the respective cells. Of note, due the absence of unique molecular identifiers in scSLAM-seq, dichotomy could also result for the over amplification of one single transcript. Our finding links dichotomous transcription to gene-specific features including TATA-box elements and CpG dinucleotides in the respective promoters. This implies that DNA methylation is associated with such non-permissive “off” states of gene promotors lasting for several hours before transcription initiation re-occurs and transcript levels are replenished. Accordingly, dichotomous transcription explains intercellular heterogeneity in cellular transcriptomes.

The early phase of CMV infection is paralleled by a strong IFN and NF-κB response, leading to the up-regulation of hundreds of genes as observed in population based measurements ^9^. Here, scSLAM-seq revealed that does not result from target genes being induced in all cells. In contrast, most of the target genes exhibit “On”-“Off” patterns in the uninfected cells, but are shifted towards more cells in the “On” state upon infection. Thus, our data indicate that the IFN and NF-κB response resolve the non-permissiveness of target promoters, leading to up-regulation at population level without increasing the maximal level of new RNA in individual cells. This may facilitate subsequent efficient pathogen recognition and control by all cells, while avoiding hyperactive cytokine responses, cell death and autoimmunity.

Metabolic labeling using 4sU is applicable to all cell types and major model organisms including vertebrates, insects, plants and yeast ^12,41^. We performed scSLAM-seq following 2hrs of metabolic labeling and found 4sU incorporation to be both efficient and uniform in all analyzed cells. 4sU incorporation into newly transcribed RNA is concentration dependent ^12^. scSLAM-seq should thus be readily applicable for 60 min of 4sU labeling or less using similar or slightly higher concentrations of 4sU. With costs for library preparation and sequencing coming down, thousands to tens of thousands of cells can be analyzed. It is compatible with other modifications of scRNA-seq including Perturb-seq ^42,43^ and will greatly improve the sensitivity of the respective approaches to decipher the molecular mechanisms underlying altered single-cell transcriptome profiles. scSLAM-seq will have major implications for developmental biology, infection and cancer.

## Acknowledgments

We thank Jörg Vogel, Wolfgang Kastenmüller and Georg Gasteiger for critical comments on the manuscript. The work was funded by the European Research Council (ERC-2016-CoG 721016-HERPES to LD). The Helmholtz Institute for RNA-based Infection Research (HIRI) supported this work with a seed grant through funds from the Bavarian Ministry of Economic Affairs and Media, Energy and Technology (Grant allocation nos. 0703/68674/5/2017 and 0703/89374/3/2017).

## Author contributions

Conceptualization: A.-E.S., F.E. and L.D.; Methodology: C.J. and F.E.; Investigation: M.A.P.B., T.K., T.H., A.P. and A.-E.S.; Writing: A.-E.S., F.E. and L.D.; Funding acquisition: A.-E.S., F.E. and L.D.; Supervision: A.-E.S., F.E. and L.D.

## Author Information

A patent has been filed on the GRAND-SLAM approach to analyze the relative contribution of transcriptional activity based on U- to C-conversions. The authors declare no other competing interests. Correspondence and request for materials should be addressed to.

## Extended data

### Extended Data Figures

1. scSLAM-seq quality controls.
2. Half-life estimates and PCA on regulated genes.
3. Correlation of viral content with incoming RNA.
4. Differentially expressed genes in CMV infected cells
5. D-scores reflect stochastic transcriptional activity in single cells
6. Sequence logos overrepresented in promoter regions of dichotomous genes

### Supplementary Information

Supplementary Tables 1 - 8.

Supplementary Methods

## Methods

### Cell culture, virus infection and RNA metabolic labeling

Murine NIH-3T3 fibroblasts were cultured in Dulbecco’s modified Eagle’s medium (DMEM) supplemented with 10 % newborn calf serum (NCS) and Penicillin Streptomycin. 24h after seeding, cells were infected with MCMV Smith strain at an MOI of 10 using centrifugal enhancement (30min at 800g). Immediately after spinocculation, virus inocculum was removed and replaced with fresh, pre-warmed medium containing 500 μM 4-thiouridine (4sU). Metabolic labeling was performed for 2hr at 37°C. Cells were detached using Trypsin/EDTA, washed twice in cold PBS and resuspended in ice cold PBS. Single cells were immediately sorted using a FACSaria III (BD Biosciences; precision: single-cell; nozzle: 100 μm) into individual wells of a 48-well plate (Brand) filled with 4 μl of a hypotonic lysis buffer containing 2 U/μL RNase inhibitor (Takara) and 0.2 % Triton-X-100 in Nuclease-free water. Single-sorted cells were spun down, immediately chilled to 4°C and stored at −80 °C. In parallel, cells from a single well of a 6-well dish (1×10^6^ cells) per condition and replicate were subjected to the same procedure (mock and CMV infection) and subjected to standard SLAM-seq as described ^7^. In this case, total cellular RNA was isolated using Trizol (Invitrogen). Two biological replicates were performed.

### Single-cell SLAM-seq

After thawing the plates for 3 min at room temperature, 0.4 μL of 10X PBS (pH=7.4, Ambion) was added to each well followed by dispensing 4.4 μL of 2X Iodoacetamide (IAA) (Thermo Scientific) to reach a final concentration of 10 mM IAA and 50% DMSO. The reaction mixture was incubated for 5 min at 50°C. The IAA reaction was quenched by adding 1.3 μL of 0.1 M DTT (final concentration of 15 mM) and incubation for 5 min at room temperature. ERCC spike-in control (Thermo Fisher, mix 1, dilution 1:2 millions) was added to each well (except for the negative control). RNA was isolated using a 1:1 volume of RNA XP magnetic beads (Beckman Coulter). After 10 min incubation at room temperature, the beads were washed twice with 50 μl 80% freshly-prepared EtOH on the magnetic stand and the dried beads pellets were resuspended in 4 μL elution buffer (RNAse inhibitor 1/200 Takara, CDS primer 0.5 μL in RNAse free water) immediately followed by an incubation at 72°C for 3 min. All the following experimental steps were performed using the Smart-seq v4 low input kit (Takara) with one-quarter of the recommended reagent volumes. Libraries were prepared using Nextera XT (Illumina) using a quarter of the recommended reagent volumes, pooled and sequenced in paired-end mode on the NextSeq500 sequencer (IIIumina) using the High Output 2×150 cycle kit.

### GRAND-SLAM 2.0

Our original GRAND-SLAM method (REF) is based on the following binomial mixture model:

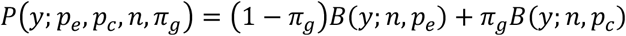

*y* is the number of T to C mismatches observed for a read covering *n* thymines, *p_e_* and *p_c_* are the probabilities of observing a T to C mismatch in RNA synthesized prior to or after 4sU labeling, respectively, and *B* is the binomial probability mass function. *p_e_* and *p_c_* can be estimated directly from the data in each individual sample. *π_g_* is the new-to-total ratio that can be estimated for each gene *g* based on the observed reads using Bayesian inference.For scSLAM-seq, we implemented the following improvements:

- SNPs are filtered by a more stringent (and statistically solid) approach: Assuming a maximal *p_c_*, a position with k substitutions in n reads is identified as a SNP by testing the Null hypothesis that *p* ≤ *p_c_* against the alternative hypothesis that *p* > *p_c_* using a Binomial test on *k~Binom*(*n*, *p*).
- For the Illumina TruSeq libraries (where the antisense strand is sequenced), A to G mismatches must be counted and incorporated into the model
- For the SMART-seq libraries (strand-unspecific sequencing), either T to C mismatches (sense reads) or A to G mismatches (antisense reads) must be counted and incorporated into the model. The strand was determined based on the gene annotations.
- For paired-end libraries (both TrueSeq and SMART-seq), the binomial model is adapted to consider read pairs instead of reads (while keeping track of them being sense or antisense).
- Many read pairs overlap, i.e. contain doubly sequenced base pairs. If the respective mismatch is observed twice, sequencing errors (but not other causes for mismatches such as PCR error, transcription errors or promiscuous RNA editing) can virtually be excluded. This is incorporated into the model by two additional parameters, *d_e_* and *d_c_*, corresponding to mismatches in doubly sequenced parts of read pairs. These can be estimated from the data using the same approach as for *p_e_* and *p_c_*.

### Data processing and analysis

Data were analyzed using the following three steps: (1) Read mapping to generate bam files of the sequencing data, (2) GRAND-SLAM analysis to produce gene tables from the bam files, (3) Downstream analyses on the obtained data. ^11^A more detailed description of the data analysis can be found in the Supplementary Methods.

Base calling was done by the internal software of the NextSeq 500 sequencer “NextSeq Control/RTA v2” and bcl2fastq2 Conversion Software v.2 was used to demultiplex the pooled libraries and to convert the BCL files generated by the sequencer to standard FASTQ files for downstream analysis. Reads were trimmed using Trimmomatic 0.36 ^44^. (Smart-seq libraries: CTGTCTCTTATACACATCTCCGAGCCCACGAGA and CTGTCTCTTATACACATCTGACGCTGCCGACGA, TruSeq Libraries: AGATCGGAAGAGCACACGTCTGAACTCCAGTCA and AGATCGGAAGAGCGTCGTGTAGGGAAAGAGTGT).

Trimmed reads were mapped to a combined reference of the mouse (Ensembl v90) and MCMV (BAC-derived Smith Strain: C3X-m129-repaired) genomes and the ERCC spike-in sequences using STAR 2.5.3a with the following parameters: --outFilterMismatchNmax 20 --outFilterScoreMinOverLread 0.4 -- outFilterMatchNminOverLread 0.4 --alignEndsType EndToEnd --outSAMattributes nM MD NH.

We first analyzed the bulk sequencing (TruSeq) libraries. Using GRAND-SLAM’s standard parameters ^11^, SNPs and the error rate regression model were inferred from all 5 samples and the no4sU bulk sample, respectively. Due to higher and more irregular error rates at the beginning of the sequencing reads, we excluded the first 10 bases from analysis by using the -trim5p 10 parameter. For the Smart-seq libraries, we first analyzed the data from the first plate which also contained the no4sU samples. We supplied the identified SNPs from the bulk sequencing experiment (by using the.snpdata file produced by GRAND-SLAM from the bulk data). Here, we ignored the first 25 bases (-trim5p 25). The identified linear regression model parameters were subsequently used to process the second plate (which did not contain any no4sU samples) via the parameters -errlm ‘TA*0.8359’ -errlm2 ‘TA*2.802’.

TPM values were computed using the read counts and mRNA lengths provided by GRAND-SLAM. RNA half-lives for the bulk samples were computed from the new/total posterior means p (estimated by GRAND-SLAM) using the formula −2log(2)/log(1-p) ^11^. Cells with less than 2,500 detected genes (TPM>1) were removed from further analyses (remaining: n=94).

Highly variable genes were identified using the ERCC spike-ins to model technical noise ^45^. We used all genes with a Benjamini-Hochberg corrected p value < 1%. The principal component analysis (PCA) for Fig. 1 was computed using the prcomp function in R. This was applied on a matrix containing either log2(1+TPM*f) (with a constant f=1 for total RNA, the new/total maximum a posteriori f=p for new RNA, or f=1-p for old RNA) or the new/total maximum a posteriori values for all cellular, highly variable genes in all 94 single cells.

Differential expression was computed with either the SCDE or the SC2P package ^18,19^. For SCDE (which performs its own normalization), we supplied the raw counts scaled with the factor f defined above and used a cutoff on the absolute value of the corrected zscore of 1.645 (corresponding to a false discovery rate (FDR) of 10%) to call differentially expressed genes. For SC2P, we first normalized reads using the sum factors approach implemented in the scran package and called genes with an FDR of 10% for both „phases“. For gene set analyses, we used gene ontology terms available via the org.Mm.eg.db R package. For transcription factor targets, we downloaded the predicted locations of transcription factor binding from the Swissregulon database ^22^ (mm10_sites_v2.gff.gz), and intersected all genomic locations with potential promotors (500 upstream to 100 downstream of the annotated transcription start site) of all transcripts from Ensembl version 90 to define sets of target genes for each of the 679 transcription factors. The PAGODA algorithm ^21^ from the SCDE package was used to identify variable gene sets based on the variance models from the differential expression analysis and while correcting for potential batch effects (based on the replicates). We then identified differential gene sets as the subset of the variable gene sets where the derived signature was significantly different for Mock and MCMV cells (based on a t test and using a Benjamini-Hochberg corrected p value cut-off of 1%). We merged all signatures from differential gene sets derived using total, new and old RNA for hierarchical clustering and merge clusters after manual inspection at dendrogram height 8 or 11 (for transcription factor target sets and GO, respectively) to average the signatures from highly similar gene sets (full data available in Supplementary Tables 4-5).

### IFN- and NF-κB-regulated genes

To investigate IFN and NF-κB targets, we downloaded the relevant target lists from MSigDB (name: HINATA_NFKB_TARGETS_FIBROBLAST_UP) or the Supplementary Table from ^12^. For CMV specific targets, we removed genes from this list that had less than 10 reads from new RNA in all four bulk samples, had an average log2 fold change in the two bulk replicates (new RNA) of less than 0.5, or were detected in less than 20 single cells (TPM>1). First, we obtained the new or old normalized (scran package) read counts (c) for those genes and computed z scores across all cells for the logarithmized counts (log2(1+c), scale function). We defined the IFN or NF-κB signature as the loadings of the first principal component performed on the cells.

### Dichotomy analysis

For the dichotomy analysis, we first selected genes where the new/total ratio was precisely quantified (90% credible interval according the beta approximation of GRAND-SLAM < 0.2) in at least 8 uninfected cells. For each gene, we utilized all precise new/total ratios across uninfected cells, and computed their standard deviation (D-score). To analyze the influence of the cell cycle, we obtained the marker genes for S and G2M phases from the Seurat package ^46^. We used the same new/total ratios as for the D-scores, discarded all new/total ratios between 0.2 and 0.8, and constructed, for each cell, a contingency table for genes using the variables “cell cycle phase” and “On or Off” (NTR>0.8 or NTR<0.2). From this contingency tables, log odds ratios and p values were computed using the fisher.test function in R. For the top 10% genes according to the D-score, we computed the correlation of all observed NTRs to the order of cells according to the log odds ratios. This procedure for 100 random permutations of this order.

Gene ontology analyses on the D-scores was performed using GOrilla ^31^. For the promoter analyses, we used the transcription factor target associations introduced above, and computed the Wilcoxon rank-sum test for targets vs. non-targets along the D-scores of each gene. P values were corrected using the Benjamini-Hochberg procedure.

### Promoter analysis

We first determined for each gene in the mouse genome its major isoform. This was done by maximizing for the annotated transcripts the number of reads starting in the first 50 nucleotides (nt) of the transcript. We obtained the position weight matrices (PWMs) from the ElemeNT database ^47^ and computed the match score at the given position in the promoter of the major isoform from each gene (we allowed for a tolerance of +/-3 nt by maximizing the score over these positions). For each PWM, we used the top 10% of all genes as “hits”, and then computed a Wilcoxon rank-sum test for the hits vs. non-hits of each PWM along the D-score of all genes. P values were corrected using the Benjamini-Hochberg procedure.

### CpG methylation

For the analysis of CpG methylation, we downloaded the processed data table for mouse fibroblasts experiment from ^38^. Genomic coordinates were transferred from mm9 to mm10 using the NCBI liftover tool. CpG island annotations were obtained from the UCSC genome browser for the mm10 genome build. For each of the major isoforms defined above, we consider the +/-500 basepair range around the annotated transcription start site. We determine whether there was an overlap with an annotated CpG island to define CGI promoters. Also within this range, we counted the number of CpGs sequenced in the bisulfite sequencing experiment, and the number of methylated CpGs. We defined a promoter as non-methylated if <10% of the CpGs was methylated.

### Data availability

The sequencing data are available at Gene Expression Omnibus (GEO) at GSE115612. The GRAND-SLAM analysis pipeline is freely available for non-commercial use at http://software.erhard-lab.de. Script files to generate all figures are available at zenodo (doi: 10.5281/zenodo.1299120).

## Extended Data Figures

**Extended Data Figure 1. scSLAM-seq quality controls.**

**a**, Total number of genes detected after scSLAM-seq across all four experimental conditions (mock- and CMV-infected cells; two biological replicates) versus the total read counts per single-cell. The horizontal line indicates a threshold below which cells have been excluded from the analysis. **b**, Partition of reads devoted for the host (cellular), the viral, the spike-in control ERCC and the mitochondrial mapped genes (Mitoch) across all the single-cells. **c**, Fraction of nucleotide substitutions per cell focused on the range 0 to 0.004 across all single-cells and 4sU-naive cells. **d**, Correlation between expression levels of bulk RNA-seq with the pooled single-cell RNA-seq data for total, new and old RNA. Genes are colored according to RNA half-life. Pearson’s correlation coefficient (R) and the number of genes used (n) are indicated. **e**, Identification of highly variable genes (HGVs) using ERCC spike-ins to model the technical noise applied to total RNA (1% false discovery rate). Squared coefficients of variation (CV^2^) are plotted against the average normalized read counts for all cells that pass the quality control filters. The solid pink line fits the average values for ERCC spike-ins (blue dots) ^45^ (solid blue curve). **f**, Principal component analysis (the two first components are depicted) of HVGs including viral genes transcriptomes. **g**, Correlation of the percentage of viral reads with the distance to uninfected cells in the first two principal components as measured by logistic regression for the PCA in panel g. **h**, The same analysis as in **g** for the PCAs in Fig. 1e.

**Extended Data Figure 2. Half-life estimates and PCA on regulated genes.**

**a**, RNA half-lives estimated from bulk SLAM-seq replicate A are scattered against estimates from replicate B. **b**, The log_2_ fold change of total RNA from bulk SLAM-seq is scattered against the log_2_ fold change of new RNA from bulk sequencing, stratified for different RNA half-lives. **c**, PCAs on genes that are differentially expressed in new RNA from the bulk experiments (absolute log_2_ fold change > 0.5). Top: PCA on genes with short RNA half-lives (<2h) are shown. Bottom: The same for long RNA half-lives (>4h). Left to Right: PCA was performed using total, old or new RNA, or the NTR. (ARI: Adjusted Rand index). **d**, Correlation analysis of the PCAs from (C) with viral reads (see Extended Fig. 1g).

**Extended Data Figure 3. Correlation of viral content with incoming RNA.**

The percentage of viral reads per cell is scattered against the total amount of old viral RNA.

**Extended Data Figure 4. Differentially expressed genes in CMV infected cells.**

Pie chart representing the number of the differentially expressed genes between mock- and CMV-infected cells and coverage between the different classes of RNA depending on their RNA-labeling status. Differentially expressed genes were called using SCDE (corrected z-score of 1.645 corresponding to a 10% false discovery rate).

**Extended Data Figure 5. D-scores reflect stochastic transcriptional activity in single cells.**

**a**, Comparison of D-score with bulk RNA expression levels stratified by RNA half-life. Shown are the r^2^ values of ordinary linear regression (lines indicated). Especially for genes with short-lived transcripts, there was no correlation indicating that high D-scores are not due to inefficient RNA capture (TPM: transcripts per million). **b**, Heatmap showing the NTR values for marker genes of the S and G2/M cell cycle phases. Grey fields indicate undetected genes. Cells (columns) were ordered according to the log odds (on-off vs. S-G2/M). Log odds and associated p values (two-sided Fisher’s exact test, corrected by Benjamini-Hochberg) are indicated on the top. Genes (rows) were ordered according to the correlation of their NTR values with the log odds ordering. **c**, Distribution of Pearson’s correlation coefficient of the top 10% most variable genes (D-score against cell cycle log odds order, see **b**). For the sake of comparison, cells were permuted randomly 100 times, and the corresponding distributions of the correlation coefficient are indicated. **d**, Correlation of RNA half-lives with the fraction of “On” genes among all genes that were either “On” or “Off”.

**Extended Data Figure 6. Sequence logos overrepresented in promoter regions of dichotomous genes.**

Sequence logos of the transcription factors with significantly enriched binding sites among genes with low (*Tbp*) or high (*Patz1, Pml, Chd1, Sin3a, Zbtb14*) D-scores obtained from the Swissregulon database.

## Supplementary Tables

**Supplementary Table 1. RNA half-lives based on new/total RNA ratios (NTR) of 10^5^ cells.**

RNA half-lives were determined for >13,500 cellular genes based on the NTR measured in a large population of cells in parallel to the scSLAM-seq experiments (two biological replicates). In addition, the virus-induced fold-changes (logFC) in total, new and old RNA are shown.

**Supplementary Table 2. List of differentially expressed genes in CMV infection.**

Differentially expressed genes during CMV infection called using single-cell two-phase testing procedure (SC2P) (10% false discovery rate) or SCDE based on either total, new or old RNA. Regulated genes in new RNA are classified by SC2P according to the mode of regulation (On/Up, Off/Down). “Corrected” z-scores as reported by SCDE are shown. In addition, the RNA half-life of the respective genes is shown.

**Supplementary Table 3. List of transcription factors (TF) identified to be involved in CMV-induced differential gene expression.**

Transcription factors with predicted binding sites (Swissregulon DB) enriched among differentially expressed genes were called using the PAGODA algorithm. They were clustered according to the similarity of their profiles across the cells, and a combined label was assigned as indicated.

**Supplementary Table 4. Gene ontologies overrepresented in cellular differentially regulated in CMV infection.**

Gene ontology terms enriched among differentially expressed genes were called using the PAGODA algorithm. They were clustered according to the similarity of their profiles across the cells, and a combined label was assigned as indicated.

**Supplementary Table 5. Signatures of IFN- and NF-κB-responsive genes in new and old RNA.**

IFN and NF-κB regulated genes in CMV infection were defined by intersecting experimental data of IFN or NF-κB induction with up-regulated genes in the SLAM-seq experiments. PCA based signatures are indicated based on old and new RNA.

**Supplementary Table 6. List of expression levels, RNA half-life and D-score for >5,500 cellular genes.**

D-scores, computed for genes where the new/total ratio was precisely quantified in at least 8 cells are indicated here in addition to the corresponding RNA half-lives and expression levels.

**Supplementary Table 7. Gene ontologies over-represented in genes with high or low D-scores.**

Over-represented GO terms were identified using GOrilla (N is the total number of genes, B is the total number of genes associated with a specific GO term, n is the number of genes in the top of the D-score sorted list and b is the number of genes in the intersection; Enrichment = (b/n) / (B/N))

**Supplementary Table 8. List of all transcription factors used for D-score related promoter analyses.**

The number of predicted targets per transcription factor (according to Swissregulon DB), the median D-scores for targets and non-targets, the p value (two-sided Wilcoxon rank-sum test) and the corrected p value (q value; Benjamini-Hochberg) are indicated.

