## Supplementary Methods for "scSLAM-seq reveals core features of transcription dynamics in single cells"

### Extended methods

Murine NIH-3T3 fibroblasts were cultured and passaged in Dulbecco's modified Eagle's medium (DMEM) supplemented with 10 % newborn calf serum (NCS) and Penicillin Streptomycin. 1x$10^5$ cells per well were seeded overnight on a 24-well plate. Cells were infected 24h later with BAC-derived wild-type MCMV Smith strain (m129 repaired) at an MOI of 10 using centrifugal enhancement at 800xg for 30min at room temperature. Mock infected cells were also subjected to spin-occulation. Immediately after spinocculation, virus inocculum was removed and replaced with fresh, pre-warmed medium containing 500 µM 4-thiouridine (4sU), which was derived from 50 mM stock solutions dissolved in water and stored at -20°C in small aliquots for one-time use. Metabolic labeling was performed for 2hr at 37°C. Cells were detached using Trypsin/EDTA and washed twice in cold PBS after centrifugation at 1,400 rpm and finally resuspended in 700 µL cold PBS.

Single cells were immediately sorted using a FACSaria III (BD Biosciences; precision: single-cell; nozzle: 100 µm) into individual wells of a 48-well plate (Brand) filled with 4 µl of a hypotonic lysis buffer containing 2 U/µL RNase inhibitor (Takara) and 0.2 % Triton-X-100 in Nuclease-free water. Single-sorted cells were spun down, immediately chilled to 4°C and stored at −80 °C. In parallel, cells from a single well of a 6-well dish (1x$10^6$ cells) per condition and replicate were subjected to the same procedure (mock and CMV infection) and subjected to standard SLAM-seq as described (*1*). In this case, total cellular RNA was isolated using Trizol (Invitrogen). Two biological replicates were performed.

***Single-cell SLAM-seq***

After thawing the plates for 3 min at room temperature, 0.4 µL of 10X PBS (pH=7.4, Ambion) was added to each well followed by dispensing 4.4 µL of 2X Iodoacetamide (IAA) (Thermo Scientific) to reach a final concentration of 10 mM IAA and 50% DMSO. The reaction mixture was incubated for 5 min at 50°C. The IAA reaction was quenched by adding 1.3 µL of 0.1 M

DTT (final concentration of 15 mM) and incubation for 5 min at room temperature. ERCC spike-in control (Thermo Fisher, mix 1, dilution 1:2 millions) was added to each well (except for the negative control). RNA was isolated using a 1:1 volume of RNA XP magnetic beads (Beckman Coulter). After 10 min incubation at room temperature, the beads were washed twice with 50 µl 80% freshly-prepared EtOH on the magnetic stand and the dried beads pellets were resuspended in 4 µL elution buffer (RNAse inhibitor 1/200 Takara, CDS primer 0.5 µL in RNAse free water) immediately followed by an incubation at 72°C for 3 min. All the following experimental steps were performed using the Smart-seq v4 low input kit (Takara) with one-quarter of the recommended reagent volumes. The PCR amplification was performed according to the manual using 24 cycles. Libraries were quantified by Qubit™ 3.0 Fluometer (ThermoFisher) and quality was checked using 2100 Bioanalyzer with High Sensitivity DNA kit (Agilent).

***Library preparation and sequencing***

0.5 ng of each library was subjected to a tagmentation-based protocol (Nextera XT, Illumina) using a quarter of the recommended reagent volumes, 10 min for tagmentation at 55 °C and 1 min extension time during PCR for multiplexing. After PCR, the libraries were purified using AMPure XP beads and eluted in 15 µl of resuspension buffer. Libraries were pooled and sequenced in paired-end mode on the NextSeq500 sequencer (Illumina) using the High Output 2×150 cycle kit.

**Data availability, processing and analysis**

Data were analyzed using the following three steps:

1. Read mapping to generate bam files of the sequencing data
2. GRAND-SLAM analysis to produce gene tables from the bam files
3. Downstream analyses on the obtained data

The sequencing data as well as the gene tables are available from GEO (GSE115612), the script files to reproduce all three steps are available at zenodo (doi: 10.5281/zenodo.1299120). GRAND-SLAM (*2*) for the second step is available for non-commercial use at http://software.erhard-lab.de.

**Step 1: Read mapping**

Base calling was done by the internal software of the NextSeq 500 sequencer "NextSeq Control/RTA v2" and bcl2fastq2 Conversion Software v.2 was used to demultiplex the pooled libraries and to convert the BCL files generated by the sequencer to standard FASTQ files for downstream analysis.

Reads were trimmed using Trimmomatic 0.36 (*3*)
(Smart-seq libraries: CTGTCTCTTATACACATCTCCGAGCCCACGAGA and
CTGTCTCTTATACACATCTGACGCTGCCGACGA,
TruSeq Libraries: AGATCGGAAGAGCACACGTCTGAACTCCAGTCA and
AGATCGGAAGAGCGTCGTGTAGGGAAAGAGTGT).
Trimmed reads were mapped to a combined reference of the mouse (Ensembl v90) and MCMV (BAC-derived Smith Strain: C3X-m129-repaired) genomes and the ERCC spike-in sequences using STAR 2.5.3a with the following parameters:
--outFilterMismatchNmax 20 --outFilterScoreMinOverLread 0.4 --outFilterMatchNminOverLread 0.4 --alignEndsType EndToEnd --outSAMattributes nM MD NH.

**Step2: GRAND-SLAM analysis pipeline**

We first analyzed the bulk sequencing (TruSeq) libraries using GRAND-SLAM and genome indices prepared from both the mouse genome (Ensembl version 90) and the MCMV genome. Using GRAND-SLAM's standard parameters, SNPs and the error rate regression model were inferred from all 5 samples and the no4sU bulk sample, respectively. Due to higher and more irregular error rates at the beginning of the sequencing reads, we excluded the first 10 bases from analysis by using the -trim5p 10 parameter. For the Smart-seq libraries, we first analyzed the data from the first plate which also contained the no4sU samples. We supplied the identified SNPs from the bulk sequencing experiment (by using the.snpdata file produced by GRAND-SLAM from the bulk data). Here, we ignored the first 25 bases (-trim5p 25). The identified linear regression model parameters were subsequently used to process the second plate (which did not contain any no4sU samples) via the parameters -errlm 'TA*0.8359' -errlm2 'TA*2.802'.

**GRAND-SLAM 2.0**

Our original GRAND-SLAM method (*2*) is based on the following binomial mixture model:

$$P(y; p_e, p_c, n, \pi_g) = (1 - \pi_g)B(y; n, p_e) + \pi_g B(y; n, p_c)$$

$y$ is the number of T to C mismatches observed for a read covering $n$ thymines, $p_e$ and $p_c$ are the probabilities of observing a T to C mismatch in RNA synthesized prior to or after 4sU labeling, respectively, and $B$ is the binomial probability mass function. $p_e$ and $p_c$ can be estimated directly from the data in each individual sample. $\pi_g$ is the new-to-total ratio that can be estimated for each gene $g$ based on the observed reads using Bayesian inference.For scSLAM-seq, we implemented the following improvements:

- SNPs are filtered by a more stringent (and statistically solid) approach: Assuming a maximal $p_c$, a position with k substitutions in n reads is identified as a SNP by testing the Null hypothesis that $p \leq p_c$ against the alternative hypothesis that $p > p_c$ using a Binomial test on $k \sim Binom(n, p)$.
- For the Illumina TruSeq libraries (where the antisense strand is sequenced), A to G mismatches must be counted and incorporated into the model
- For the SMART-seq libraries (strand-unspecific sequencing), either T to C mismatches (sense reads) or A to G mismatches (antisense reads) must be counted and incorporated into the model. The strand was determined based on the gene annotations.
- For paired-end libraries (both TrueSeq and SMART-seq), the binomial model is adapted to consider read pairs instead of reads (while keeping track of them being sense or antisense).
- Many read pairs overlap, i.e. contain doubly sequenced base pairs. If the respective mismatch is observed twice, sequencing errors (but not other causes for mismatches

such as PCR error, transcription errors or promiscuous RNA editing) can virtually be excluded. This is incorporated into the model by two additional parameters, $d_e$ and $d_c$, corresponding to mismatches in doubly sequenced parts of read pairs. These can be estimated from the data using the same approach as for $p_e$ and $p_c$.

**Step 3: Downstream analyses**

TPM values were computed using the read counts and mRNA lengths provided by GRAND-SLAM. RNA half-lives for the bulk samples were computed from the new/total posterior means p (estimated by GRAND-SLAM) using the formula -2log(2)/log(1-p) (*2*). Cells with less than 2,500 detected genes (TPM>1) were removed from further analyses (remaining: n=94).

Highly variable genes were identified using the ERCC spike-ins to model technical noise (*4*). We used all genes with a Benjamini-Hochberg corrected p value < 1%. The principal component analysis (PCA) for Fig. 1 was computed using the prcomp function in R. This was applied on a matrix containing either log2(1+TPM*f) (with a constant f=1 for total RNA, the new/total maximum a posteriori f=p for new RNA, or f=1-p for old RNA) or the new/total maximum a posteriori values for all cellular, highly variable genes in all 94 single cells. For each PCA, we also computed a logistic regression model for the Infection status (Mock/MCMV) based on the first two principal components. We calculated the area under the receiver operating characteristics curve statistics for each regression model using the pROC package for R.

For Fig. S2, we used the same procedures, but computed PCAs not on highly variable genes, but on genes that were at least 2x induced or 2x repressed in the bulk sequencing experiment (new RNA). Expression fold changes for the bulk sequencing experiments were computed using PsiLFC (*5*). Adjusted Rand indices were computed using the clues R package.

For the analysis of MCMV genes, we utilized the data from (*6*) and defined early genes as the ones measured before 6.5h post infection (p.i.), late genes as the ones measured at first at

24 h.p.i. and true late genes as the ones measured at first at 48 h.p.i. For each gene, we considered $\log_2(1+TPM)$ values and normalized the corresponding estimates for old and new RNA to the maximal value for total RNA across all cells.
