## Supplementary figures and images for "scSLAM-seq reveals core features of transcription dynamics in single cells"

### Supplementary Figures

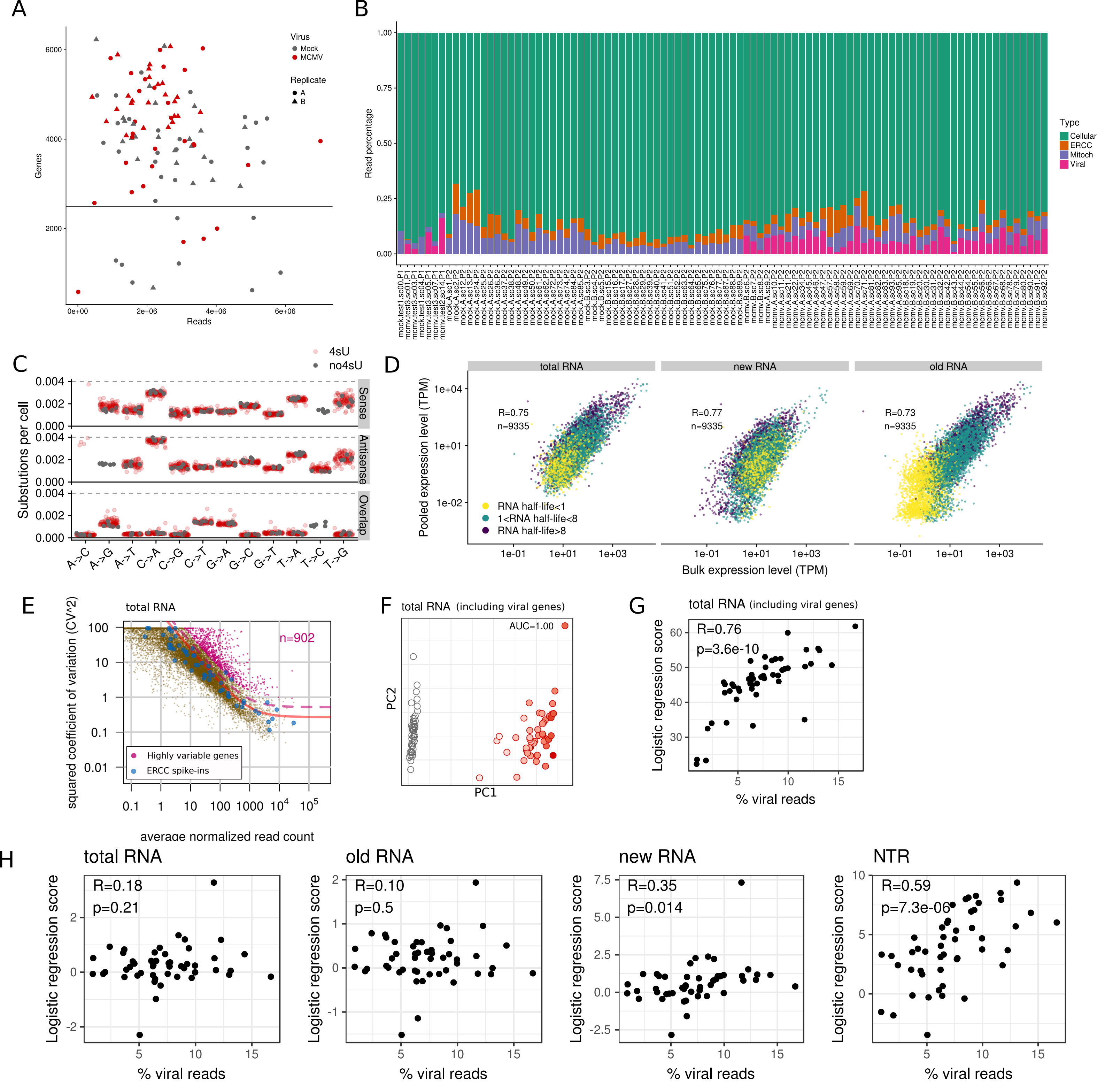

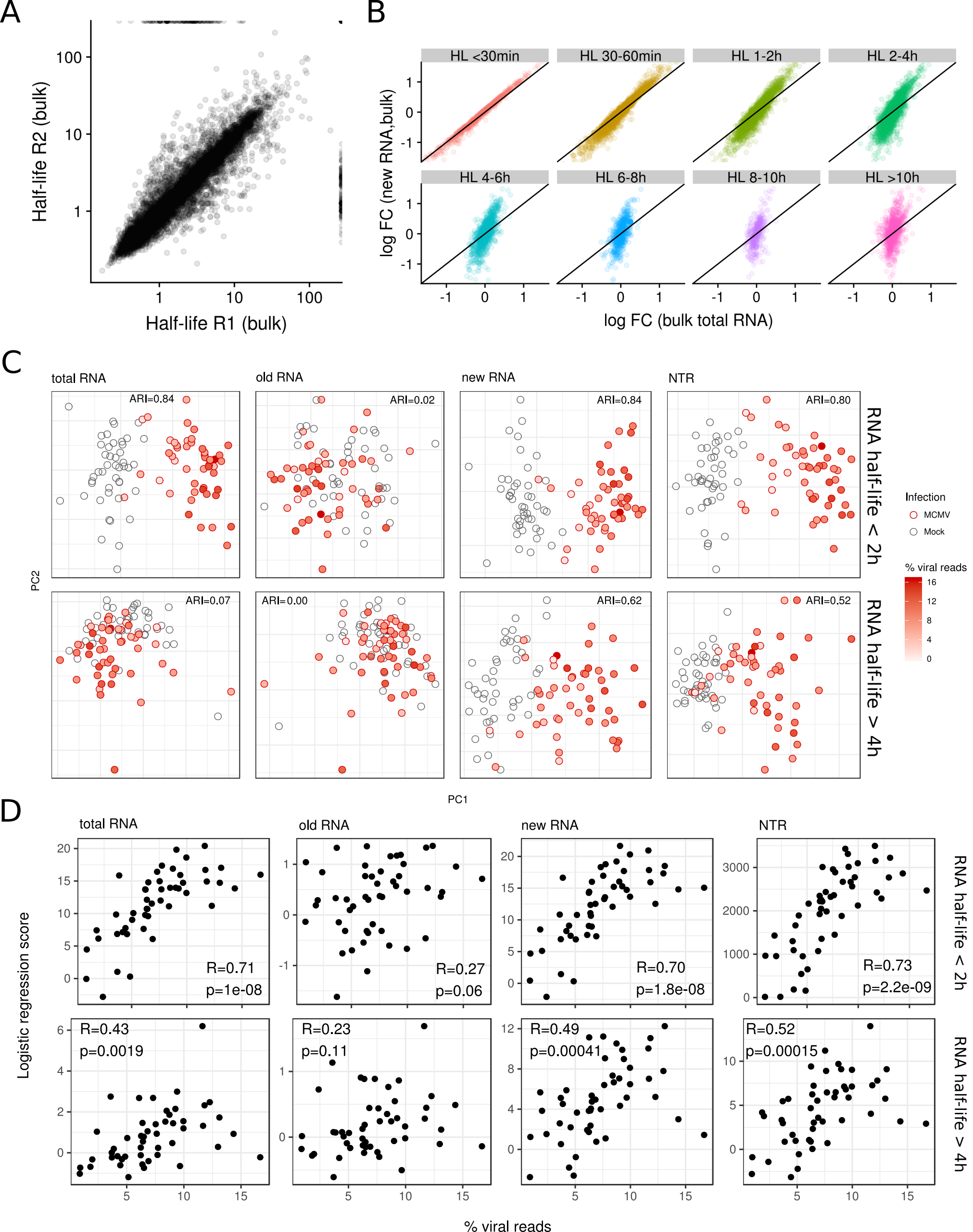

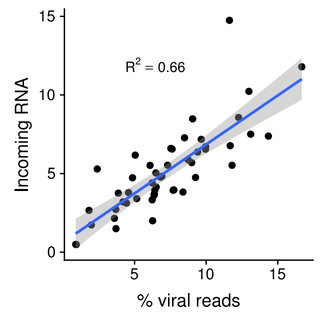

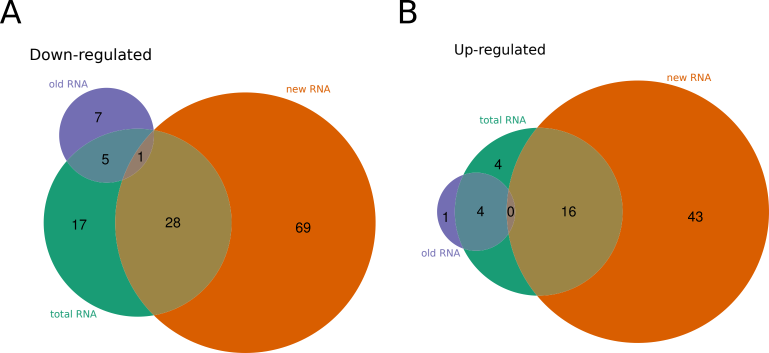

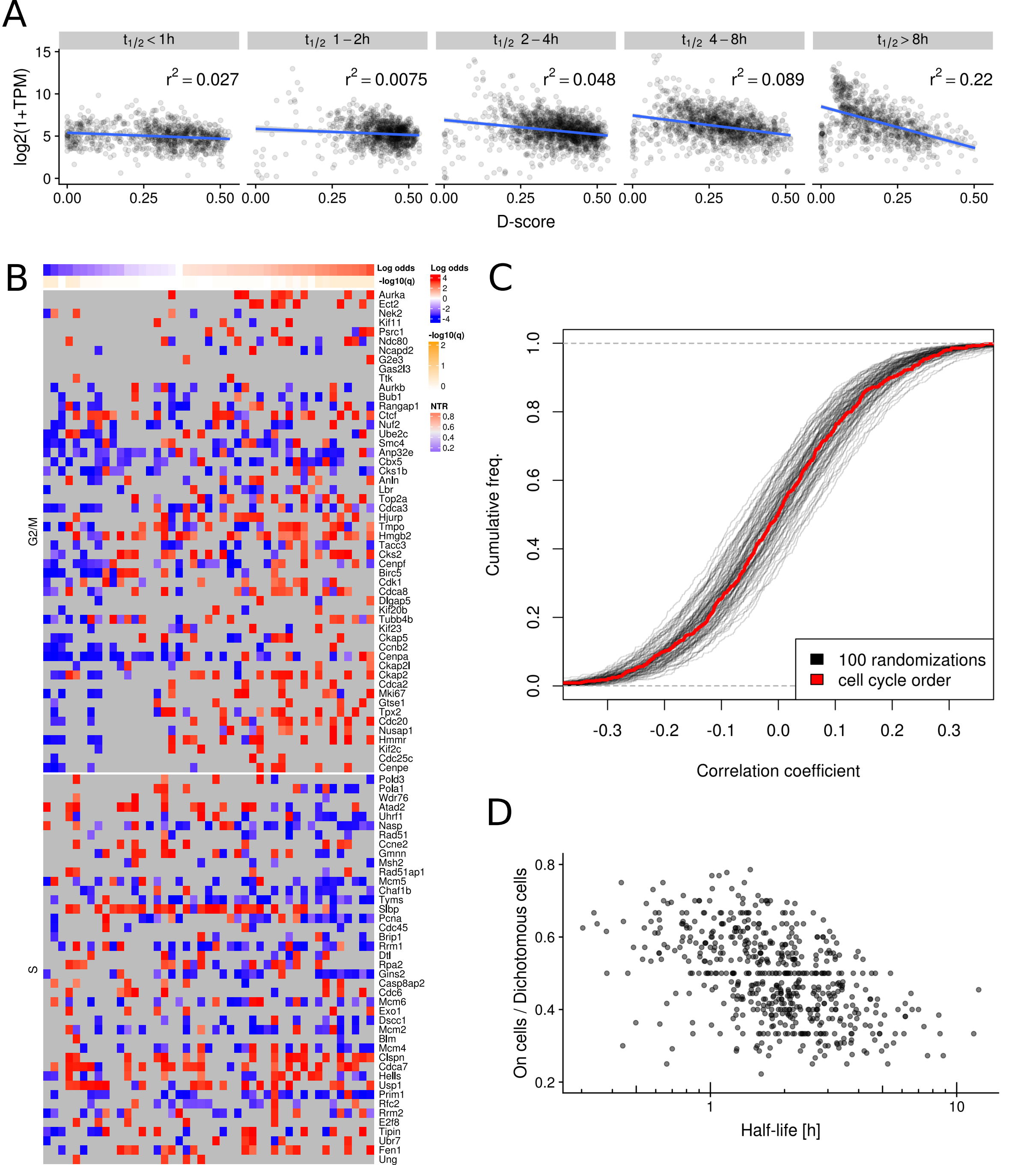

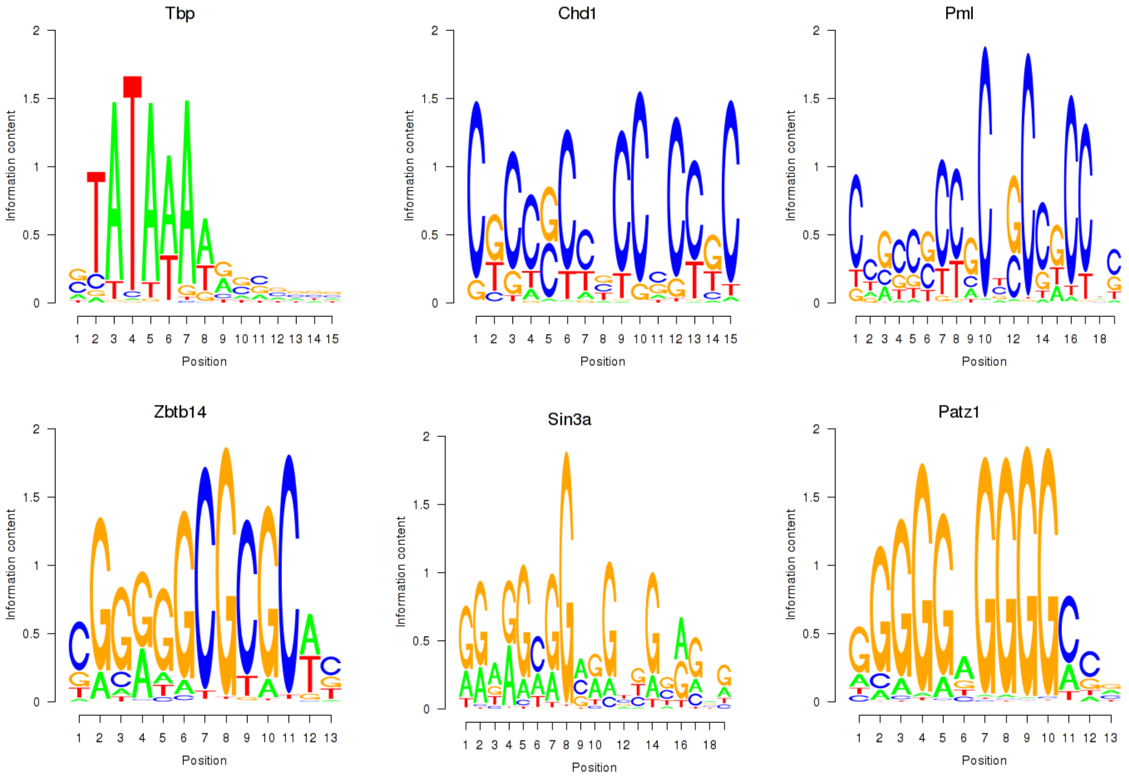
